# Altered Motor Development Following Late Gestational Alcohol and Cannabinoid Exposure in Rats

**DOI:** 10.1101/513713

**Authors:** Kristen R. Breit, Brandonn Zamudio, Jennifer D. Thomas

## Abstract

Cannabis is the most commonly used illicit drug among pregnant women, and rates are likely to increase given recent legalization. In addition, half of pregnant women who report consuming cannabis also report drinking alcohol. However, little is known about the consequences of prenatal cannabis alone or combination with alcohol, particularly with cannabis products that are continually increasing in potency of the primary psychoactive constituent in cannabis, Δ9-tetrahydrocannabinol (THC). The current study investigated the effects of early exposure to cannabinoids during the brain growth spurt on early physical and motor development alone (Experiment 1) or in combination with alcohol (Experiment 2). In Experiment 1, Sprague-Dawley rat pups were exposed to a cannabinoid receptor agonist (CP-55,940 [CP]; 0.1, 0.25, 0.4 mg/kg/day), the drug vehicle, or a saline control from postnatal days (PD) 4-9. In Experiment 2, rat pups were exposed to CP (0.4 mg/kg/day) or the vehicle, and were additionally intubated with alcohol (11.9% v/v; 5.25 g/kg/day) or received a sham intubation. Subjects in both experiments were tested on a motor development task (PD 12-20) and a motor coordination task during adolescence (PD 30-32). Both developmental cannabinoid and alcohol exposure separately decreased body growth throughout development, and combined exposure exacerbated these effects, although only alcohol exposure induced long-term body weight reductions. Developmental cannabinoid exposure advanced early motor development, whereas alcohol exposure delayed development, and subjects given combined exposure did not differ from controls on some measures. Alcohol exposure impaired motor coordination later in life. In contrast, cannabinoid exposure, by itself did not significantly affect long-term motor coordination, but did exacerbate alcohol-related impairments in motor coordination among females. These results suggest that cannabinoid exposure may not only alter development by itself, but may exacerbate alcohol’s teratogenic effects in specific behavioral domains. These findings have important implications not only for individuals affected by prenatal exposure, but also for establishing public policy for women regarding cannabis use during pregnancy.

## 1. Introduction

Cannabis is the most commonly used illicit drug among women of reproductive age^1^. It is also the most commonly used illicit drug used among pregnant women^2^ in the United States, with prevalence rates ranging from 3-4%^3^ and higher rates among pregnant adolescents^4^. Given the recent legalization of cannabis in many states, the perception that cannabis is safe to consume during pregnancy^5^, and the reported intention of use among pregnant women despite knowledge of potential risks^6^, the availability and use of cannabis among pregnant women is likely to increase. Importantly, maternal ingestion of Δ9-tetrahydrocannabinol (THC), the primary psychoactive constituent in cannabis, can have direct effects on fetal development, as THC and its metabolites can freely pass across the placenta^7^. However, relatively little is known about the effects of prenatal cannabis exposure on fetal development.

There are few prospective clinical studies that have examined the effects of prenatal cannabis exposure on early development, and results from these and retrospective studies are mixed^8,9^. Prenatal cannabis exposure generally does not produce physical birth defects, although it may reduce birth weight^9–15^ and possibly alter emotional, behavioral, and cognitive development, although results have been inconsistent^9^. These inconsistencies are likely due to differences in cannabis exposure levels, prospective versus retrospective approaches, confounds of other drug use, age, and nature of outcome measures, as well as a host of other methodological, maternal, and environmental factors. Of particular concern is the continually increasing potency of cannabis-related products that are currently available compared to past levels. The potency of THC in cannabis-related products has continually risen from 3.4% to 12.2% from 1993 to 2015 and higher percentages of psychoactive compounds among synthetic cannabinoids^16,17^, with an average potency of 11% in 2016 in cannabis-related products^18^. However, it will be years before any long-term consequences of prenatal exposure to high potency cannabis are known, as THC levels have even further increased since the 2001 initiation of the most recent prospective Generation R study.

Studies using animal models similarly report mixed results of prenatal cannabis exposure on behavioral development in emotional^19^, cognitive^20^, and motor domains^21^; however, these results vary drastically based on differences in timing, dose and form of cannabinoid, outcome measures, and nature of control groups^9,22–24^. For example, prenatal cannabinoid exposure has been shown to increase spontaneous motility (ambulation and rearing) in rats as early as PD 10^25^, PD 13^26^, and PD 15 when challenged with THC^27^, and at PD 12 with the cannabinoid agonist WIN 55,212-2^28^. However, other rodent studies have shown decreased motility (THC)^29^, no changes (THC)^30,31^, and sex-dependent stereotypy motor alterations (hashish)^32^ following developmental cannabinoid exposure. Zebrafish embryos briefly exposed to THC during gastrulation exhibit altered morphology of motor neurons, neuromuscular junction synaptic activity, and locomotor responses to stimuli^33^. Similarly, prenatal THC exposure in mice may disrupt cortical connectivity in areas associated with motor development, leading to long-lasting reductions in fine motor skills^34^. However, no studies to date have provided information about initial motor development in rats following early cannabinoid exposure or how altered development may relate to later motor skills.

Importantly, cannabis is not the only drug consumed by pregnant women. A recent survey suggested that 5.5% of women of reproductive age report using cannabis and alcohol simultaneously, with up to 15.3% of women between 18-29 years of age reporting simultaneous use^35^. In fact, cannabis is the most common illicit drug used simultaneously with alcohol among women who report binge drinking during pregnancy^36^. According to the National Household Survey on Drug Abuse, approximately 50% of pregnant women who report consuming cannabis also report drinking alcohol^37^.

In contrast to the inconsistent findings regarding prenatal cannabis exposure, the dangers of alcohol consumption during pregnancy are well established. Individuals exposed to alcohol prenatally may suffer from a range of physical, neurological, and behavioral consequences referred to as fetal alcohol spectrum disorders (FASD), which may include growth deficits, and impairments in a range of cognitive and behavioral domains, including impaired motor function^38,39^. Developmental alcohol exposure has been shown to impair both fine and gross motor function in both clinical^40–42^ and preclinical studies^40,43,44^. However, it is largely unknown whether concurrent cannabis exposure may exacerbate alcohol’s teratogenic effects on early physical and motor development.

Our understanding of the effects of concurrent prenatal cannabis and alcohol exposure is severely lacking, as most clinical studies focus on the effects of each drug separately^11,45–47^, rather than the combination of effects^19^. Similarly, animal models have primarily focused only on the teratogenic effects of alcohol or cannabinoids individually; few have examined the combination, and those that did have focused on neurotoxicity^48,49^. For example, the combination of ethanol and THC exposure is neurotoxic when administered during the first 2 postnatal weeks in the rodent^48,49^; however, studies investigating the behavioral effects of combined prenatal cannabinoid and alcohol exposure are limited. Thus, the functional consequences of combined developmental exposure to alcohol and cannabis on early physical and motor development remain largely unknown. Given that exposure to either substance alone can disrupt fetal development, if the combination produces additive or synergistic effects on early development, both personal and public health costs could be extensive.

To examine the possible consequences of developmental exposure to cannabinoids as well as combined exposure to cannabinoids and alcohol on early development, we used a rat model of drug exposure during PD 4-9, a time period that corresponds to the late gestational brain growth spurt. The brain growth spurt is a period of axonal growth, dendritic arborization, high rates of synaptogenesis, gliogenesis, myelination, and maturation of synaptic neurotransmission^50,51^. The endogenous cannabinoid system also plays an important functional role in neuronal development during this period, influencing proliferation, migration and synaptogenesis^52^. CB_1_ receptor levels rapidly increase in most brain areas at this time, including the cerebral cortex, hippocampus, brainstem, septum nuclei, cerebellum, and striatum^53–55^. Importantly, this is also a period of development particularly sensitive to ethanol^56,57^.

To mimic the effects of THC, the synthetic CB_1_ receptor agonist CP-55,940 (CP) was used, as it is one of the most well-characterized and commonly used compounds in cannabinoid research^58^. Although more potent, the peak effect, duration of action, and neurobehavioral effects of CP are almost identical to those of THC^59^, and it is also the main ingredient in several “synthetic marijuana” preparations available for human use^60^. In Experiment 1, CP was administered at various doses to model low, moderate and high exposure levels of THC in human consumption^61,62^. In Experiment 2, CP was combined with alcohol administered in a binge-like manner at a dose known to produce physical and behavioral alterations^63–65^.

Body weights, developmental milestones, and blood alcohol concentrations were measured to investigate possible effects of independent and combined exposure to cannabinoids and alcohol during development. To examine early motor development, a grip strength and hindlimb coordination task^43,66^ was used from PD 12-20, a period of critical of motor and sensory development^67,68^. To examine motor coordination later in adolescence, a parallel bar motor coordination paradigm was used from PD 30-32^65^. All subjects were tested on both tasks to compare possible changes in motor performance across time following developmental cannabinoid and/or alcohol exposure.

## 2. Material and Methods

All procedures included in this study were approved by the San Diego State University (SDSU) Institutional Animal Care and Use Committee (IACUC) and are in accordance with the National Institute of Health’s Guide for the Care and Use of Laboratory Animals.

### 2.1. Experiment 1 Design

#### 2.1.1. Subjects

Sprague-Dawley rat offspring subjects were bred onsite at the SDSU Animal Care facilities at the Center for Behavioral Teratology. During breeding, a male and a female rat were housed together overnight and the presence of a seminal plug the following morning indicated gestational day (GD) 0. Dams were then individually housed, and except for routine monitoring and maintenance, remained undisturbed until GD 22, the typical day of delivery. At birth, litters were pseudo-randomly culled to 8 animals (4 sex pairs, whenever possible). Subjects were randomly assigned within a litter to each treatment group, and no more than one sex pair per litter was used for each treatment condition so that no litter was overrepresented in any treatment group.

#### 2.1.2. Developmental Cannabinoid Exposure

The purpose of Experiment 1 was to investigate the potential effects of neonatal exposure to clinically relevant low, medium, and high doses of cannabinoids on early motor development and later motor coordination. Drug exposure occurred between postnatal days (PD) 4-9, a model of the late gestational “brain growth spurt.” All subjects received intraperitoneal (i.p.) injections (10 ml/kg) of one of three doses of CP-55,940 (0.40 [CP4], 0.25 [CP2.5], 0.10 [CP1] mg/kg/day), whereas control subjects received 10% DMSO vehicle (VEH) or physiological saline (SAL). During cannabinoid exposure, all pups were removed from the dam and maintained on a heating pad; each injection took approximately one minute per pup to complete. A total of 120 subjects was used in Experiment 1.

On PD 7, offspring were tattooed for identification purposes with non-toxic tattoo ink, allowing the experimenter to remain blind during testing. All motor testing occurred from PD 12-32. On PD 21, offspring were weaned and housed together, and on PD 28, subjects were separated by sex. All animals were housed at a constant humidity and temperature, and exposed to a 12-hour light/dark cycle, receiving food and water ad libitum. All testing and procedures were conducted during the subjects’ light cycle.

#### 2.1.3. Drug Preparation

CP-55,940 (CP; Enzo Life Sciences, NY) was dissolved into a stock solution (5 mg of CP dissolved into 2mL of 100% Dimethyl Sulfoxide [DMSO] Sigma-Aldrich, MO) and kept at −20°C until daily solutions were made. Daily injection volumes were prepared by combining the CP stock solution with the vehicle (10% DMSO and physiological saline) to the appropriate final doses (0.40, 0.25, 0.10 mg/kg/day), using serial dilutions.

### 2.2. Experiment 2 Design

#### 2.2.1. Subjects

A separate cohort of subjects was generated for Experiment 2, bred at the SDSU Animal Care facilities at the Center for Behavioral Teratology as in Experiment 1.

#### 2.2.2. Developmental Alcohol and Cannabinoid Exposure

The goal of Experiment 2 was to examine the effects of combined neonatal alcohol and cannabinoid exposure on early motor development and later motor coordination. From PD 4-9, half of all subjects were intragastrically intubated with ethanol (EtOH, 5.25 g/kg/day) dissolved in an artificial milk diet (11.9% v/v, twice per day, 2 hours apart), followed by 2 additional feedings of milk diet only^69^. Briefly, flexible polyethylene-10 tubing (Intramedic, Clay Adams Brand, USA) was attached to a 25-gauge needle on a 1 mL syringe to create the intragastric intubation equipment. The tubing was lubricated with corn oil, passed over the tongue into the esophagus, and slid into the stomach. The EtOH milk or milk solution was delivered within a 10-sec period.

Remaining subjects were fully intubated with the tubing (Sham), with no EtOH exposure. In addition, all subjects were injected (i.p.10 ml/kg) with either the highest dose of CP from Experiment 1 (0.40 [CP4] mg/kg/day) or the drug vehicle (10% DMSO and physiological saline [VEH]). A total of 148 offspring subjects was used for Experiment 2. Similar to Experiment 1, all pups were removed from the dam during intubations and maintained on a heating pad; each intubation and injection took approximately one minute per pup to complete.

On PD 6, 20 microliters of blood was collected from each subject via tail clip for blood alcohol concentration (BAC) analyses (Analox Alcohol Analyzer, Model AMI; Analox Instruments; Lunenburg, MA) to examine if cannabinoid exposure altered BAC levels. Peak blood alcohol concentrations (BACs) were measured 90 minutes after the last alcohol intubation on postnatal day 6^70^. Sham-intubated subjects also had 20 microliters of blood collected to control for possible stress effects, although samples were not analyzed. Tattoos (PD 7), motor testing (PD 12-32), weaning (PD 21), and separate housing (PD 28) occurred in an identical timeline to Experiment 1.

#### 2.2.3. Drug Preparation

The EtOH (95%, Sigma-Aldrich) was added to an artificial milk diet at 11.9% v/v. The milk diet consisted of evaporated milk (Carnation^®^), soy protein isolate and vitamin diet fortification mix (MP Biomedicals, CA), corn oil (Mazo^®^), methionine, tryptophan, calcium phosphate, deoxycholate acid, and other essential minerals (Sigma-Aldrich, MO) present in rat dam lactation^71^. CP-55,940 for Experiment 2 was prepared in the same manner as in Experiment 1.

### 2.3. Early Physical Development (Experiments 1 and 2)

Milk band presence was recorded each day throughout drug administration to ensure maternal feeding during both Experiments 1 and 2. Body weights and eye opening (day when both eyes were fully open) were recorded each day from PD 4-20 (throughout exposure and the motor development testing) and body weights were also recorded on the first day of the parallel bar motor coordination test.

### 2.4. Early Motor Development (Experiments 1 and 2)

On PD 12-20, subjects were tested on a developmental grip strength and hindlimb coordination task to examine early sensorimotor maturation and motor development, as previously described^43^. Subjects were given 2 trials per day and allowed 30 seconds (sec) to complete each trial. Each pup was suspended from a wire (2-mm diameter) by the two front paws above a cage filled with bedding. A successful grip strength trial was recorded if the subject was able to hold on to the wire for 30 sec. A successful hindlimb coordination trial was recorded if the pup was able to place one of its hindlimbs on the wire within 30 sec of having its forepaws placed on the wire. If subjects achieved either a successful grip strength or hindlimb trial on the 2 trials allowed in one day, that day was recorded as a double success day. All success types were assessed during testing by an investigator, but trials were also recorded using a video camera for later reference, if needed.

### 2.5. Motor Coordination (Experiments 1 and 2)

From PD 30-32, subjects were tested on a parallel bar motor coordination paradigm to examine motor coordination during adolescence. The apparatus consists of two parallel steel rods (0.5-cm diameter, 91-cm length) held between two end platforms (15.5 x 17.8 cm) that had grooved slots spaced 0.5 cm apart to secure the rods and varying widths. The rods were 63 cm above a cage filled with bedding. Subjects were placed on one platform for 30 sec to acclimate, then placed halfway between the platforms on the parallel bars. Four alternating steps with the hindlimbs constituted a successful traversal; with each success, the distance between the bars was increased by 0.5 cm. Falling or swinging off the rods was considered an unsuccessful traversal, and subjects were allowed five unsuccessful trials at a given width before testing was terminated for the day. A maximum of 15 trial attempts per day was allowed. The number of unsuccessful attempts before the first successful trial as well as the maximum width (cm) achieved each day were recorded. The overall success ratio was calculated by dividing the number of successful trials by the total number of trials attempted.

### 2.6. Statistical Analyses (Experiments 1 and 2)

Dependent variables for physical development included body weights (grams [g]) and milk band presence (binomial) on each day, as well as the first day of eye opening (both eyes fully open). Dependent variables for the hindlimb coordination task included the age of first success, and total number of successes, and ability to succeed on each day. Data were analyzed separately for recorded grip strength trials, hindlimb coordination trials, and double success days. Dependent variables for the parallel bar motor coordination task included the number of trials to first success, maximum width achieved, and success ratios.

For Experiment 1, data that were normally distributed were analyzed using 5 (CP: CP4, CP2.5, CP1, VEH, SAL) x 2 (sex: female, male) ANOVAs. For Experiment 2, normally distributed data were analyzed using 2 (EtOH exposure: EtOH, Sham) x 2 (CP exposure: CP4, VEH) x 2 (sex: female, male) ANOVAs. Repeated measures ANOVAs for Day were used when applicable. Post hoc analyses were conducted using Student-Newman-Keuls. For both Experiments, data that were not normally distributed (Shapiro-Wilk test) were analyzed nonparametrically using Mann-Whitney, Kruskal-Wallis, or Fishers-exact analyses, where appropriate. Means (M) and standard errors of the mean (SEM) are reported when applicable. All significance levels were set as p < 0.05.

## 3. Results

### 3.1. Experiment 1: Effects of CP Exposure

A total of 11 subjects was lost during treatment (CP4: 0 female [F], 1 male [M]; CP2.5: 2 F, 1 M; CP1: 0 F, 2 M; VEH: 2 F, 1 M; SAL: 0 F, 3 M); 109 subjects completed the entire behavioral paradigm, with 10-12 subjects represented in each exposure group and sex (CP4: 12 F, 11 M; CP2.5: 11 F, 11 M; CP1: 11 F, 11 M; VEH: 11 F, 10 M; SAL: 11 F, 10 M).

#### 3.1.1. Body Weights

Although all subjects gained significant weight from PD 4-9 (p’s < 0.001; Figure 1), exposure to CP impaired body growth across treatment days (Group x Day interaction; F[20,495] = 8.8, p < 0.001). On PD 4, there were no differences in body weights among groups. However, from PD 5-9, body weights of subjects exposed to CP began to lag compared to that of controls; by the end of the treatment period (PD 9), all subjects exposed to CP, regardless of dose, weighed less than VEH and SAL controls (p’s < 0.05). Overall, females weighed less than males (F[1,99] = 13.0, p < 0.001), although Sex as a variable did not interact with either CP exposure or Day. Importantly, milk band presence did not differ based on CP exposure or Sex.

**Figure 1.**
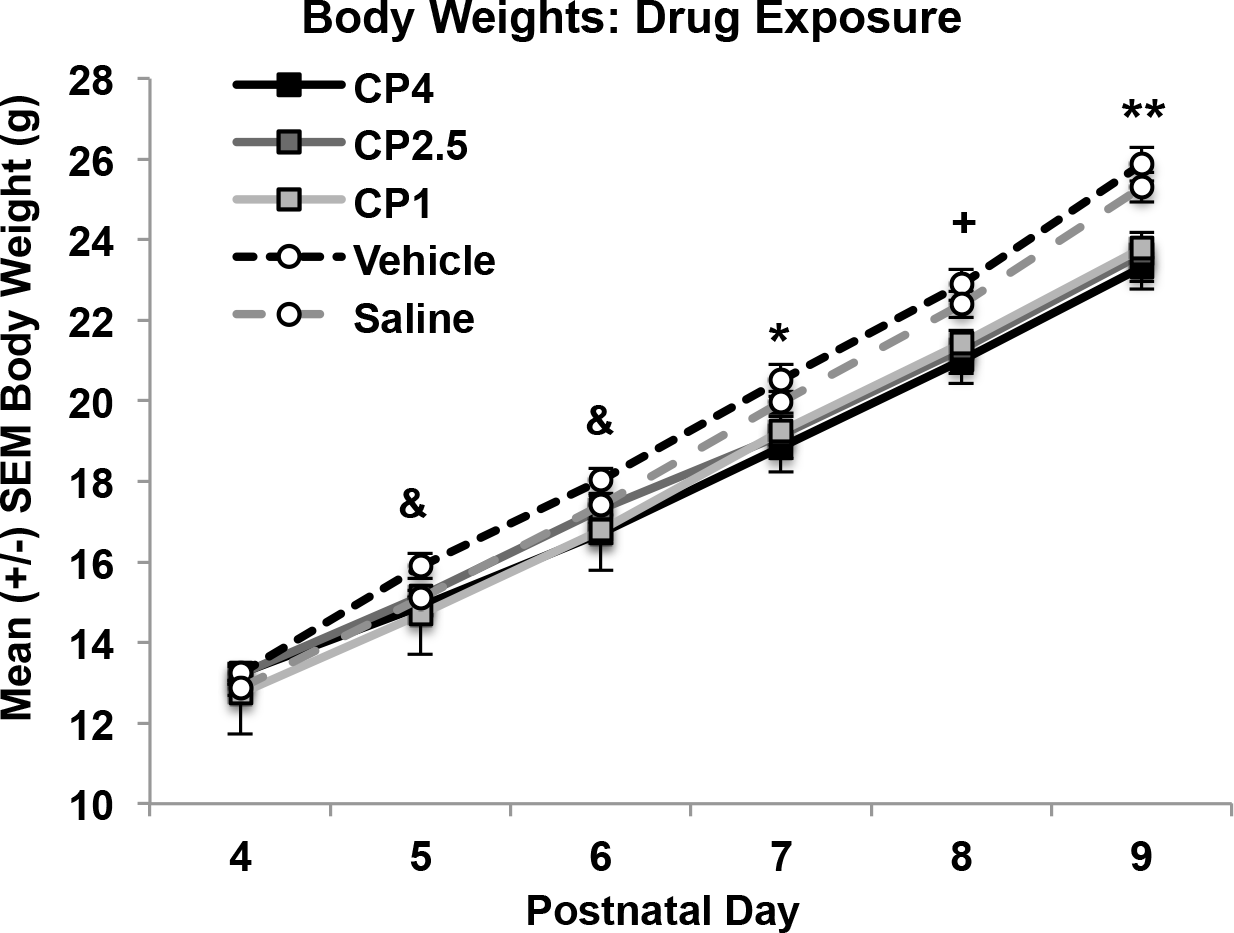
Mean (+/- SEM) body weights (grams) of subjects during drug exposure collapsed across sex. Subjects exposed to the cannabinoid receptor agonist (CP-55,940) weighed less than both control groups by the end of the exposure period. & = CP4 and CP1 < VEH; + = CP4 < VEH and SAL, CP2.5 and CP1 < VEH; * = all CP < VEH; ** = all CP < VEH and SAL.

By PD 20, subjects exposed to CP during development no longer differed in body weight from controls (body weights (g): F[4,99] = 1.56, p =.19; CP4: 54.26 ± 0.68; CP2.5: 55.26 ± 0.95; CP1: 54.81 ± 1.01; VEH: 56.98 ± 1.04; SAL: 55.51 ± 0.75). Females continued to weigh less than males on PD 20 (F[4,99] = 12.19, p < .001).

#### 3.1.2. Day of Eye Opening

Neither CP exposure (F[4,99] = 1.2, p = 0.34) nor Sex (F[1,99 = 0.7, p = 0.41) altered the first day of eye opening (CP4 [Mean ± SEM]: 13.61 ± 0.14; CP2.5: 13.91 ± 0.15; CP1: 13.77 ± 0.15; VEH: 13.53 ± 0.15; SAL: 13.58 ± 0.15).

#### 3.1.3. Early Motor Development

CP exposure did significantly improve hindlimb coordination from PD 15-20. In particular, more subjects exposed to CP4 and CP1 had achieved success compared to the controls on PD 15 and 16, whereas only subjects exposed to the highest dose (CP4) had achieved more success compared to controls from PD 17-20 (Fisher’s exact probability, p’s < 0.05; see Figure 2A). Thus, the highest dose of CP significantly advanced development of hindlimb coordination on this task. When examining overall success on each outcome measure, increases in success failed to reach significance, unless data were collapsed across CP dose, although the effect size on these measures was small. Finally, there were no significant main or interactive effects of Sex any outcome measure.

**Figure 2.**
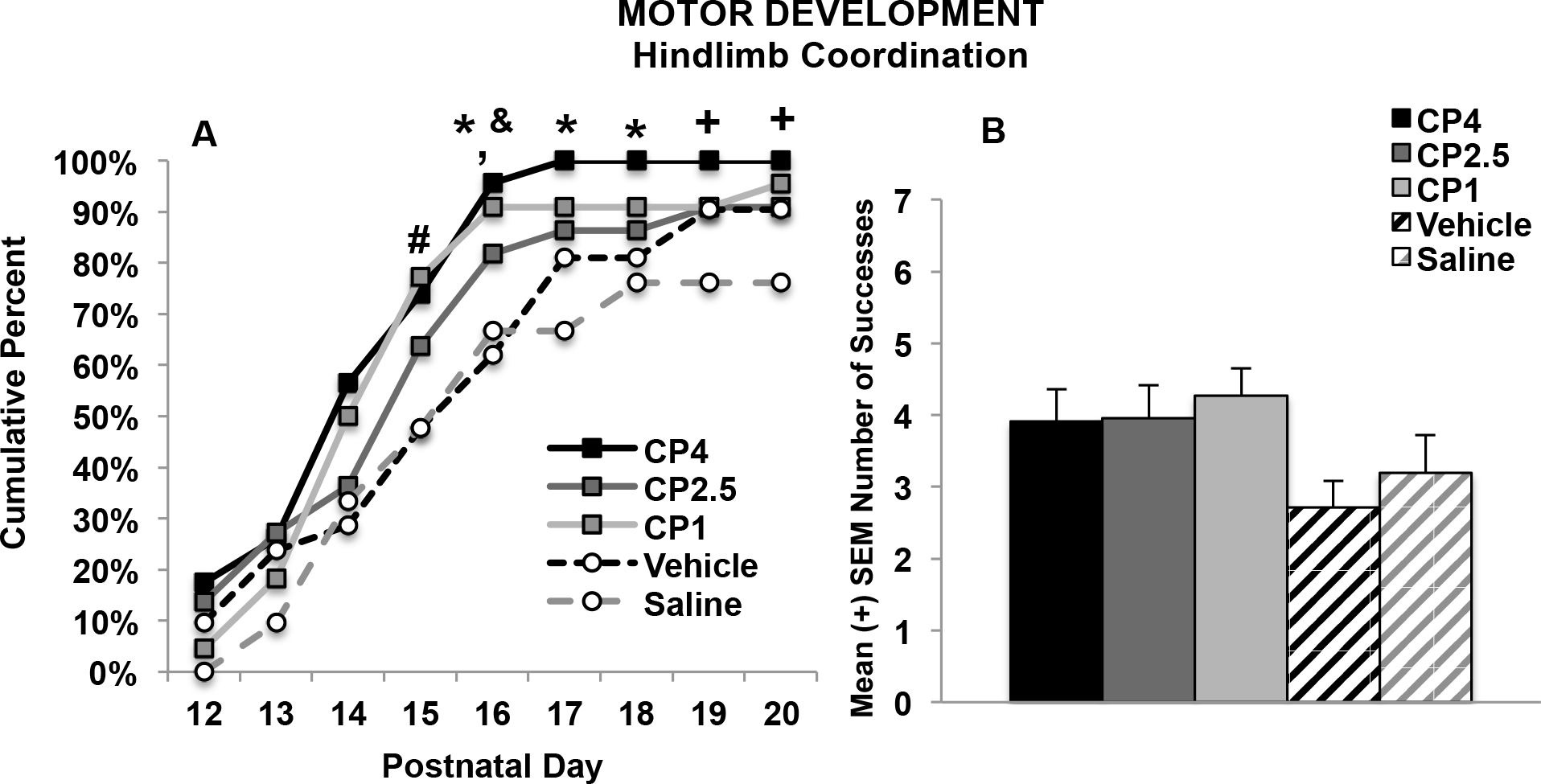
Percent of successful subjects (A) and total number of successes (B) in each exposure group that were able to place their hindlimb on the wire within 30 seconds. Subjects exposed to developmental cannabinoids, particularly the high (CP4) and low (CP1) cannabinoid doses, were able to be more successful at an earlier age (A). # = CP1> VEH and SAL; * = CP4 > VEH and SAL; & = CP1 > VEH; + = CP4 > SAL.

#### 3.1.4. Parallel Bar Motor Coordination

In contrast to the cannabinoid-related effects on motor development, there were no significant long-term effects of CP on motor coordination. There was a significant main effect of group on the mean number of trials to the first successful traversal on the parallel bars (F[1,99] = 2.63, p < 0.05; CP4: 12.78, ± 0.7, CP2.5: 13.09 ± 0.8, CP1: 10.32 ± 0.8, VEH: 10.86 ± 0.9, SAL: 10.43 ± 1.0). However, post-hoc analyses yielded no significant differences among groups. There were no effects of Sex on this measure. Similarly, there were no significant differences in the percent of subjects able to traverse the parallel bars at least once during testing. Developmental CP exposure also did not significantly affect the maximum width between bars successfully traversed on the parallel bar motor coordination task (Figure 3A). A Day*Sex interaction confirmed that males performed worse than female subjects, regardless of CP exposure (F[2,198] = 6.15, p < 0.01; data not shown), particularly on Days 2 and 3 of testing (p’s < 0.05), although both sexes improved performance across testing (p’s < 0.001). Similarly, CP exposure did not statistically significantly affect the success ratios (see Figure 3B); males performed significantly worse than females (F[1,99] = 8.14, p < 0.01; data not shown). Notably, outcome measures on this task were not significantly correlated with body weight, so sex effects were not related to body size.

**Figure 3.**
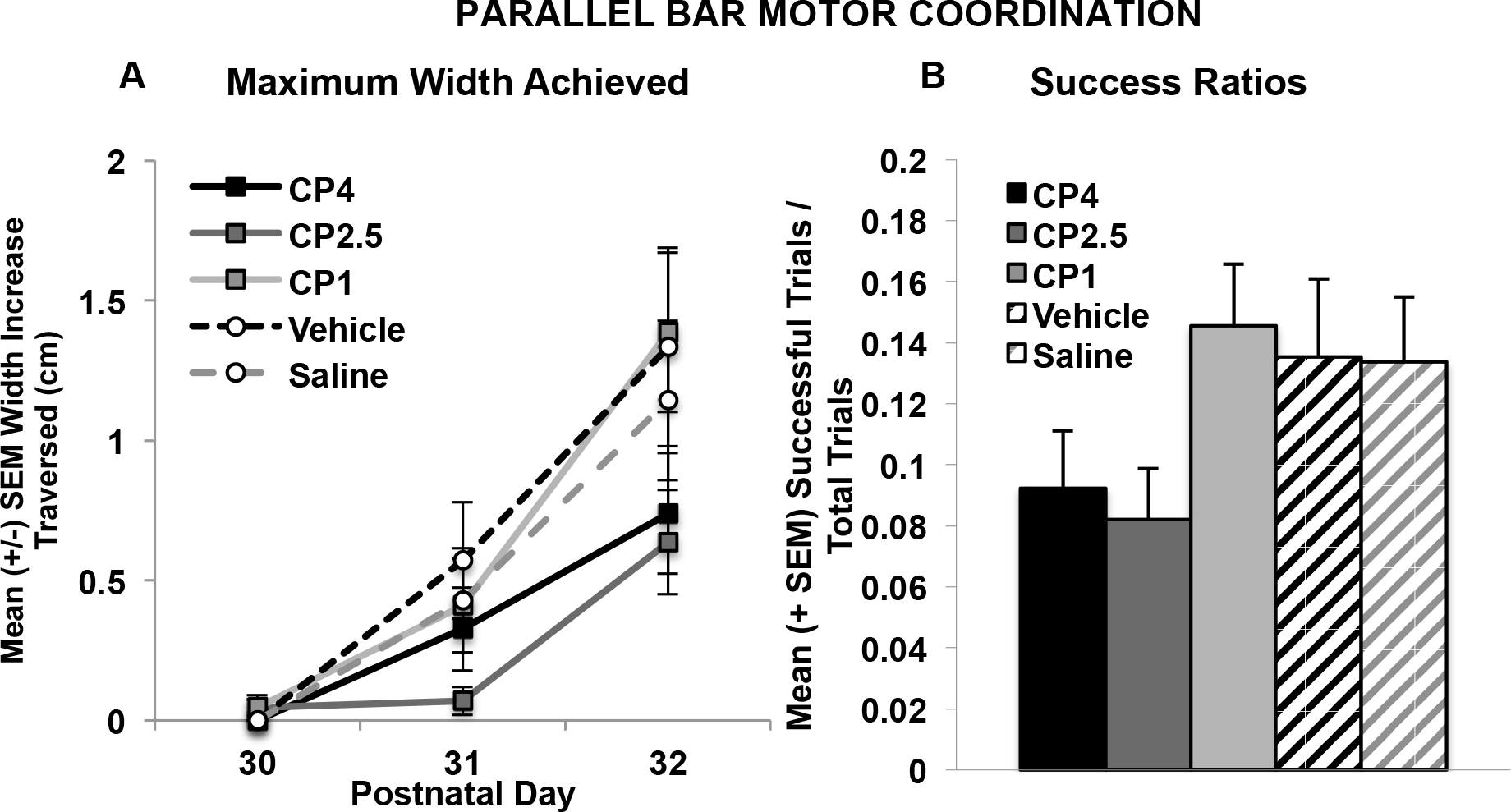
Maximum width achieved on each day (A) and motor coordination success ratios (B) on the parallel bar motor coordination test. Developmental exposure to the cannabinoid receptor agonist (CP-55,940) did not significantly affect motor performance.

### 3.2. Experiment 2: Effects of CP and Ethanol Exposure

Mortality was particularly high among subjects exposed to both ethanol and CP. A total of 31 subjects was lost due to treatment in Experiment 2 (EtOH+CP: 9 F, 11 M; EtOH+VEH: 4 F, 7 M; Sham+CP: 0 F, 0 M; Sham+VEH: 0 F, 0 M); 111 subjects completed behavioral testing in Experiment 2, with 10-16 subjects represented in each exposure group and sex (EtOH+CP: 13 F, 10 M; EtOH+VEH: 16 F, 14 M; Sham+CP: 15 F, 15 M; Sham+VEH: 13 F, 15 M).

#### 3.2.1. Body Weights

Although body weights did not differ among groups on the first day of exposure (PD 4), body growth lagged in subjects exposed to alcohol, an effect that was exacerbated with the combined exposure of ethanol and CP, producing significant interactions of Day*EtOH*CP (F[5,515] = 9.42, p < 0.01; see Figure 4A), EtOH*CP (F[1,103] = 5.13, p < 0.05), and main effects of EtOH (F[1,103] = 54.15, p < 0.001) and CP (F[1,103] = 14.06, p < 0.001). In contrast to Experiment 1, CP exposure by itself did not significantly affect body growth. Overall, females weighed less than males (F[1,103] = 7.09, p < 0.01). Similar to Experiment 1, milk band presence was not altered by EtOH exposure, CP exposure, or Sex.

**Figure 4.**
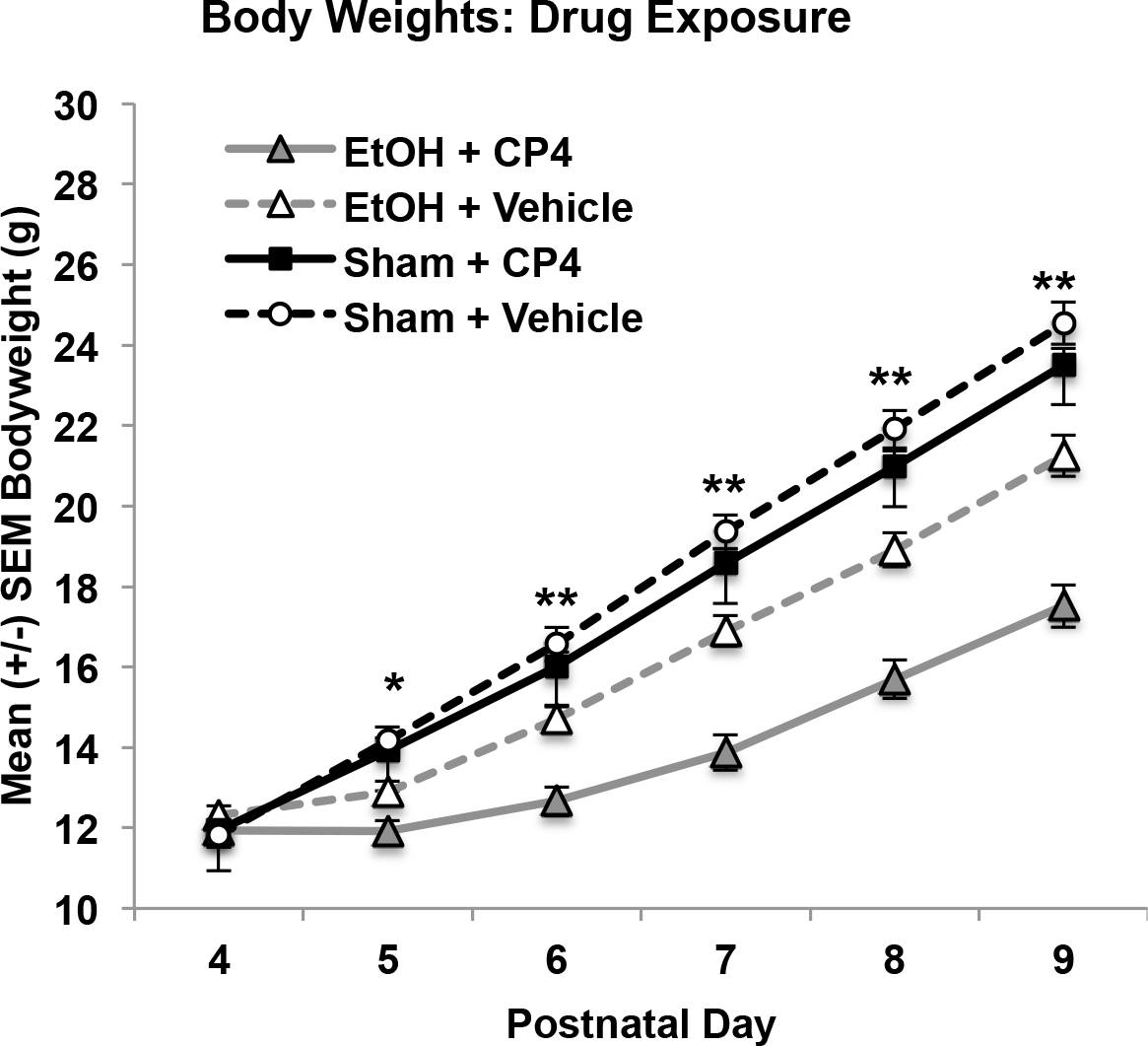
Mean (+/- SEM) body weights (grams) of subjects in during drug exposure collapsed across sex. The combination of developmental ethanol and CP-55,940 exposure decreased body weights even more than ethanol exposure alone during the treatment period * = EtOH < Sham ** = EtOH+CP < all groups and EtOH+VEH < Sham.

By PD 20, subjects exposed to EtOH during development continued to weigh less than sham-intubated subjects, producing a main effect of EtOH (F[1,103] = 39.2, p < 0.001; Figure 4B). Although the interaction of EtOH and CP failed to reach statistical significance (F[1,103] = 3.50, p = 0.06), there was a main effect of CP exposure (F[1,103] = 6.5, p < 0.05), driven by the reductions in body weight by subjects exposed to both EtOH and CP (body weights (g): EtOH+CP: 45.56 ± 1.61; EtOH+VEH: 52.38 ± 1.70; Sham+CP: 57.92 ± 1.18; Sham+VEH: 58.99 ± 1.33). Females did not weigh significantly less than males on PD 20 (F[1,103] = 3.21, p = 0.08; data not shown).

By PD 30, EtOH-exposed subjects still weighed less than Sham-intubated subjects (F[1,103] = 22.59, p < 0.001; EtOH+CP: 103.35 ± 3.57; EtOH+VEH: 110.15 ± 3.26; Sham+CP: 122.68 ± 3.25; Sham+VEH: 121.86 ± 3.19), while CP-exposed subjects no longer differed from VEH or SAL controls, and female subjects weighed less than males (F[1,103] = 25.82, p < 0.001).

#### 3.2.2. Blood Alcohol Concentrations (PD 6)

Ethanol-exposed subjects also given CP had significantly higher blood alcohol concentrations (BAC = 316.24 ± 15.15 mg/dl) than subjects not given CP (276.43 ± 13.18 mg/dl; F[1,49] = 3.93, p = 0.05). Although the interaction between CP and sex failed to reach significance, CP-related increases in BACs were driven by the females (EtOH+CP: 347.57 ± 17.09 vs. EtOH: 279.08 ± 15.41 mg/dl; F[1,27] = 8.86, p < 0.01), more than the males (EtOH+CP: 284.92 ± 26.26 vs EtOH: 273.78 ± 22.19 mg/dl; n.s.).

#### 3.2.3. Day of Eye Opening

EtOH exposure during the brain growth spurt significantly advanced the first day of eye opening (F[1,103] = 6.30, p < 0.05; EtOH: 13.63 ± 0.09; Sham: 13.94 ± 0.09). However, consistent with Experiment 1, neither CP exposure (F[1,103] = 0.26, p = 0.62) nor Sex (F[1,103] = 1.49, p = 0.23) significantly affected development of eye opening.

#### 3.2.4. Early Motor Development

Subjects exposed to CP achieved a successful grip strength trial at an earlier age than VEH-exposed subjects (F[1,103] = 3.89, p = 0.05), whereas developmental EtOH exposure did not significantly affect the first day of success (EtOH+CP: 14.61 ± 0.52; EtOH+VEH: 15.00 ± 0.38; Sham+CP: 13.47 ± 0.33; Sham+VEH: 14.82 ± 0.55). However, Fisher’s-exact analyses on each day of motor development testing indicated that subjects exposed to CP alone were more successful hanging onto the wire, whereas subjects exposed to ethanol alone were less successful, early in testing (PD 12-14, PD 16; p’s < 0.05; Figure 5A). Performance of subjects exposed to both CP and ethanol was intermediate, not differing significantly from that of controls. Overall, subjects exposed to CP during development had a greater number of successful grip strength trials compared to VEH subjects (main effect of CP: F[1,103] = 8.54, p < 0.01) (EtOH+CP: 3.70 ± 0.45; EtOH+VEH: 2.92 ± 0.33; Sham+CP: 4.47 ± 0.41; Sham+VEH: 2.96 ± 0.42).

**Figure 5.**
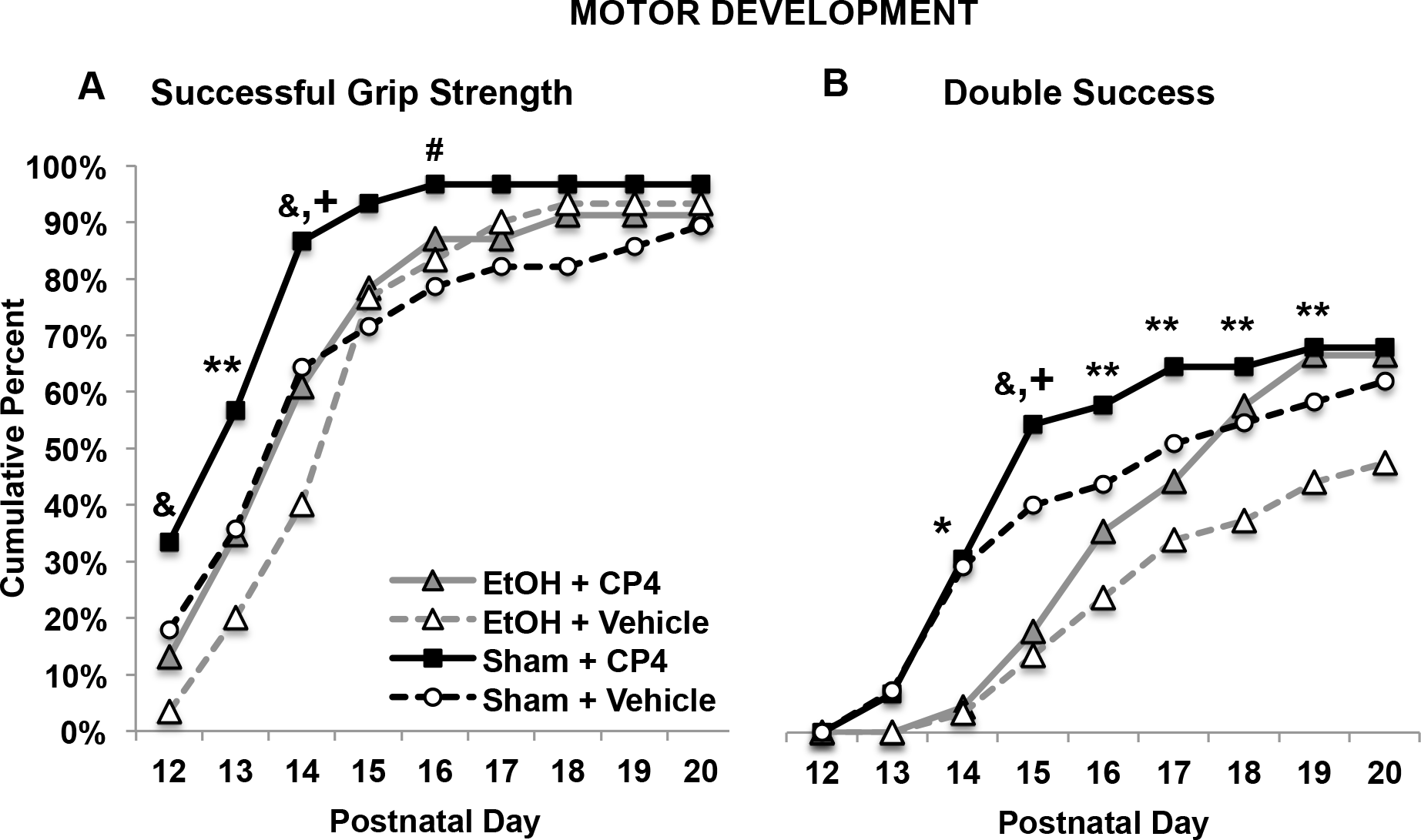
The percentage of subjects in each group able to hold onto the wire for 30 seconds (A) and that had two successful trials in one day (B) during motor development testing (collapsed across sex). A: Exposure to CP advanced the ability to perform a grip strength trial, whereas ethanol exposure delayed ability. B: Developmental ethanol exposure reduced double success days. Subjects exposed to CP were more successful, whereas subjects exposed to the combination were delayed in their successful performance, but eventually performing as well as those exposed to CP only. & = EtOH+VEH < Sham + VEH and Sham + CP; ** = Sham+CP > EtOH+VEH. + = Sham+CP > EtOH+CP. # = Sham+CP > Sham+VEH. * = Sham > EtOH.

A similar pattern of CP advancing and ethanol delaying motor development is seen with double success performance. There were no main effects of CP exposure on the first double success day; however, subjects exposed to EtOH took longer to achieve their first double success day compared to Sham subjects, producing a significant main effect of EtOH (F[1,103] = 5.05, p < 0.05; EtOH+CP: 18.07 ± 0.53; EtOH+VEH: 18.98 ± 0.45; Sham+CP: 16.93 ± 0.58; Sham+VEH: 17.59 ± 0.61). When looking at the percent of subjects that had achieved a double success by each day during motor development testing, EtOH-exposed groups were less successful by PD 14 (Fisher’s exact probability, p < 0.05). By PD 15, subjects exposed to EtOH alone performed worse than subjects exposed to CP alone and controls, whereas subjects exposed to CP alone performed better than both ethanol-treated groups (p’s < 0.05). From PD 16-19, the subjects exposed to ethanol only were only less successful than subjects exposed to CP only (p < 0.05), with performance of the combined EtOH + CP group and controls being intermediate and not significantly different from other groups (Figure 5B). When examining the overall total number of double success days, there was a main effect of ethanol (F[1,103] = 4.17, p < 0.05) and a main effect of CP (F[1,103] = 4.74, p < 0.05), as ethanol reduced success and CP increased success (EtOH+CP: 1.33 ± 0.36; EtOH+VEH: 0.70 ± 0.20; Sham+CP: 2.03 ± 0.37; Sham+VEH: 1.34 ± 0.31).

Finally, neither CP nor EtOH exposure during a model of late gestation altered the first day of hindlimb coordination success (EtOH+CP: 16.09 ± 0.60; EtOH+VEH: 16.83 ± 0.50; Sham+CP: 16.40 ± 0.55; Sham+VEH: 17.50 ± 0.59). In contrast with Experiment 1, there were no significant differences among groups in percent of subjects successful each day of testing. However, consistent with Experiment 1, subjects exposed to CP during development had a greater number of hindlimb coordination successes over testing than VEH subjects, producing a main effect of CP (F[1,103] = 6.18, p < 0.05; EtOH+CP: 2.74 ± 0.47; EtOH+VEH: 2.00 ± 0.32; Sham+CP: 2.97 ± 0.42; Sham+VEH: 1.73 ± 0.36). There were no differences between male and female subjects for any motor development measures.

#### 3.2.5. Parallel Bar Motor Coordination

In contrast to motor development, CP exposure did not improve performance and, in fact, exacerbated ethanol-related impairments on some measures. First, subjects exposed to EtOH during development required more attempts before completing a successful trial on the parallel bar task (F[1,103] = 27.42, p < 0.01), regardless of CP exposure (EtOH+CP: 14.94 ± 0.5; EtOH+VEH: 13.57 ± 0.6; Sham+CP: 11.10 ± 0.8; Sham+VEH: 10.11 ± 0.7). Neither CP exposure nor Sex significantly affected this measure.

Similar to Experiment 1, CP did not significantly affect the maximum width between rods traversed. However, subjects exposed to EtOH during development showed little improvement over training, performing worse than Sham-intubated subjects on Days 2 and 3 of testing (p’s < 0.01; Figure 6), regardless of CP exposure, and producing a Day*EtOH interaction (F[2,206] = 41.46, p < 0.001) and main effect of EtOH (F[1,103] = 35.35, p < 0.001). There were no differences between male and female subjects in the maximum width (cm) achieved.

**Figure 6.**
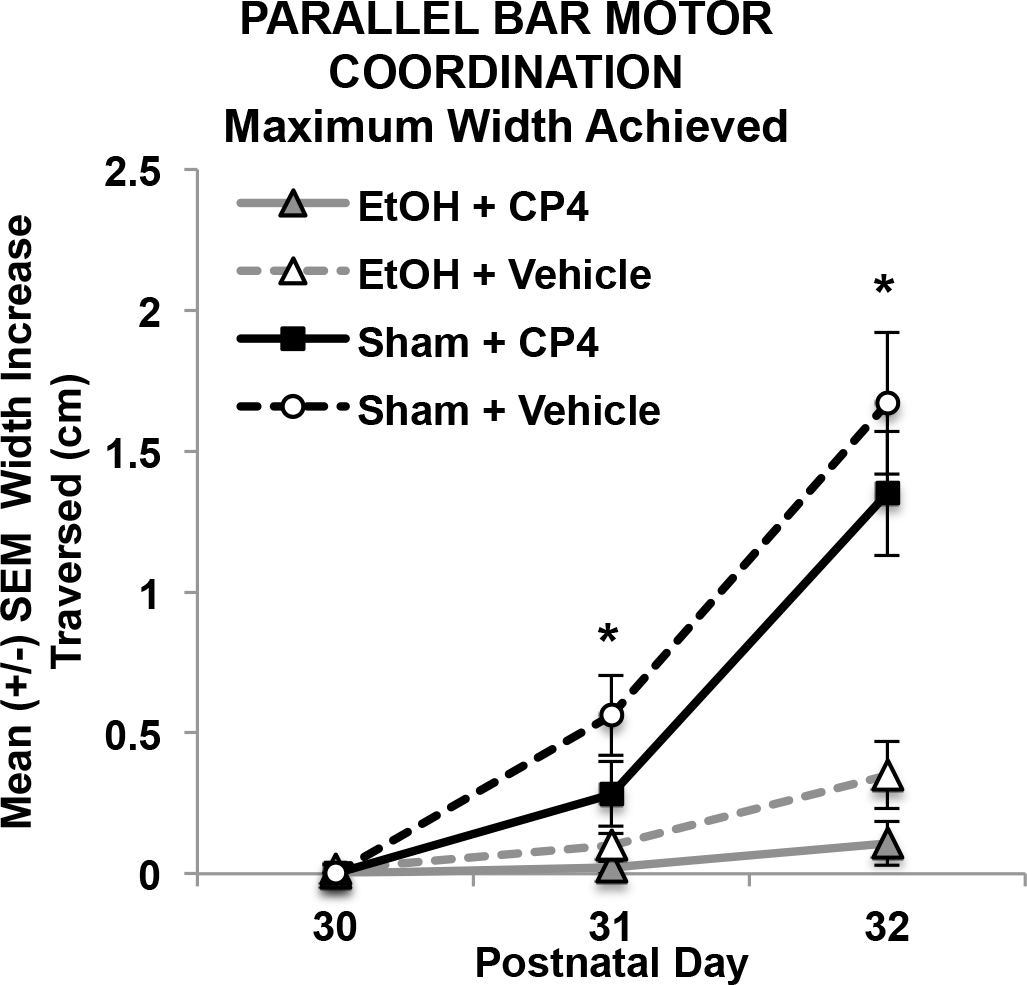
Maximum width (cm) increase achieved on each day of parallel bar motor coordination testing subjects (collapsed across sex). Ethanol-exposed subjects showed little improvement in performance over days. * = EtOH<Sham.

However, a 3-way interaction of Sex*EtOH*CP was observed in the success ratios of subjects (F[1,103] = 4.67, p < 0.05; Figure 7). Ethanol exposure impaired motor performance in both sexes, but female subjects exposed to the combination of EtOH+CP during development were less successful than female subjects exposed to EtOH alone (F[1,27] = 8.28, p < 0.01), whereas CP by itself did not significantly alter performance. Among male subjects, EtOH exposure decreased subjects’ success ratios (F[1,50] = 28.99, p < 0.001), but CP exposure did not significantly alter performance in either the EtOH- or Sham-intubated subgroups. However, it is important to note that possible interactive effects in male subjects may have been masked by a floor effect in the EtOH-exposed males. Additionally, there were no significant correlations between body weight and any outcome measures on the parallel bar motor coordination task.

**Figure 7.**
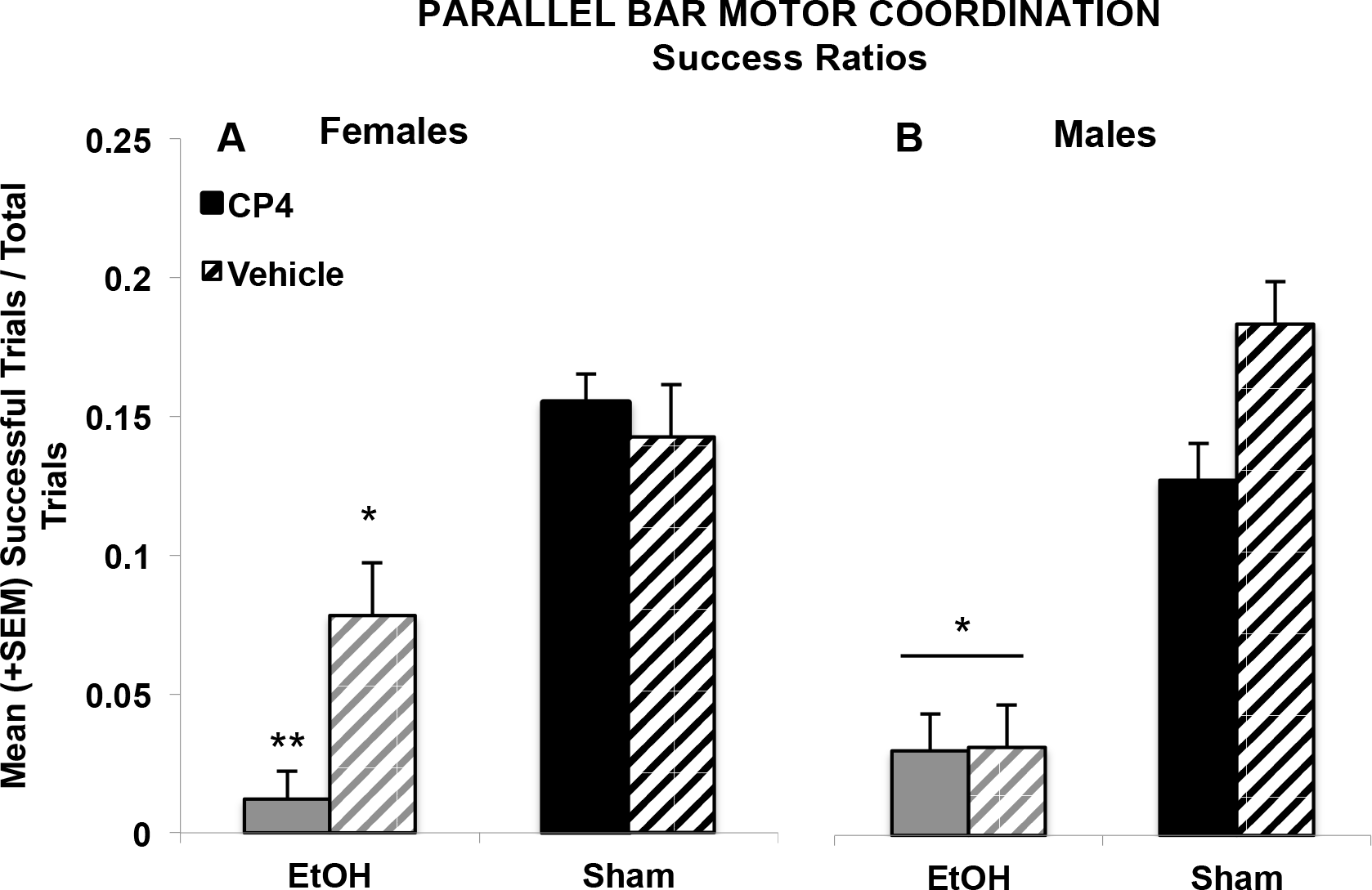
Developmental ethanol exposure impaired parallel bar success ratios in both female and male subjects; however, combined ethanol and CP exposure further impaired success ratios in female subjects more than ethanol exposure alone. * = EtOH<Sham; ** = EtOH+CP< all other female groups

## 4. Discussion

This study demonstrated that exposure to a cannabinoid during a portion of the human 3^rd^ trimester equivalent of central nervous system maturity can alter motor development and that the combination of cannabinoid and alcohol exposure during this period of development led to unique additive and synergistic effects. Developmental CP exposure advanced early motor development (as measured by grip strength and hindlimb coordination), whereas developmental ethanol delayed motor development, and the combination led to performance that reflected opposing directional effects, not differing from controls on some measures, but delaying followed by faster catch-up on other measures. Ethanol-related impairments in motor coordination and balance persisted into adolescence, whereas CP had no long-lasting effects on motor performance. However, the CP exposure did exacerbate ethanol-related motor impairments among female subjects, suggesting that the combination of a cannabinoid and ethanol during early development may produce more severe long-lasting behavioral deficits on some tasks.

Physical growth was also affected by developmental exposure to these drugs. Consistent with previous studies^63,65,72^, developmental EtOH exposure slowed body growth. The effects of CP exposure alone on body growth were inconsistent, with lags in growth observed in Experiment 1, but not Experiment 2. Importantly, only ethanol exposure had long-lasting effects on body weight by adolescence (PD 30). Few studies have administered cannabinoids during the brain growth spurt period, and no studies to date have examined combined exposure and its effects on physical development. However, a similar effect has been observed in previous prenatal cannabinoid studies. Prenatal THC exposure in rats has been shown to inhibit offspring body growth during their first 5-7 days of life, followed by a growth spurt, with normalization between PD 11-32, dependent on THC dose^73,74^.

In past studies, inhibited body growth of cannabinoid-exposed offspring has been attributed to possible alterations in the neonatal environment, such as poor nursing or cannabinoid absorption through lactation (see review by ^22^); this association has been supported by cross-fostering studies^75^. However, in the current study, both cannabinoid and ethanol exposure occurred postnatally, mimicking late gestational exposure (PD 4-9); thus, all dams were naïve to drug exposure, thereby avoiding maternal nursing-related confounds. Importantly, each litter contained subjects from all treatment groups and there were no overt differences in maternal behavior towards pups assigned to the different treatment groups in either experiment; we did videotape home cage activity and observed no obvious differences in maternal behavior (data not shown). In addition, no noticeable differences were observed in milk band presence of pups.

As expected, developmental ethanol exposure generally impaired performance in the grip strength and hindlimb coordination task, a finding previously shown by our laboratory using a prenatal alcohol model^43,66^. Ethanol-related motor impairments persisted into adolescence, as ethanol-exposed subjects were less successful on all measures of the parallel bar motor task. In contrast, developmental CP exposure advanced the developmental trajectory of motor coordination maturation in measures of grip strength and hindlimb coordination. Subjects exposed to CP during development were generally more successful on these motor outcomes and were able to perform them at an earlier age; however, it is important to reiterate that these results were only seen in Experiment 1 when collapsed across CP dose, due to small effect sizes. To our knowledge, this is the first preclinical study to show that developmental cannabinoid exposure may improve early motor performance. Importantly, behavioral testing began 3 days after the last CP exposure. Thus, due to the shorter half-life of CP^76^ when compared to substances such as THC^77^, it is unlikely that the behavioral alterations observed among CP-exposed subjects were due to the continued presence of the drug, but rather related to changes in development. Moreover, a previous study by Fried and Watkinson (1990) suggested that superior motor performance was detected in 36-month-old children born to moderate (> 1 joint/week < 6 joints/week) marijuana users^47^. Children were assessed using the motor scale in the McCarthy Scales of Children’s Abilities^78^, which includes measures for both upper- and lower-body components, and the period of assessment corresponded roughly to the developmental period we assessed in the present study (PD 12-20).

During this time of early development, preclinical studies have shown high levels of CB_1_ receptor expression in GABAergic neurons in the rat cerebellum^79^, a brain area noted for its role in motor coordination^80^. For example, within the cerebellum, cannabinoid receptor binding capacity nearly doubles from PD 0 to PD 7 and doubles again from PD 7 to 14. Thus, it is possible that the advancement in motor development seen in the current study may be partly due to CP activating cannabinoid type-1 (CB1) receptors early during development and altering cerebellar GABA neurotransmission, although our data would suggest that any changes did not persist into adulthood. Indeed, prenatal exposure to WIN 55,212-2, another CB_1_ receptor agonist, is associated with long-lasting up-regulation of GABA immunoreactivity^81^, and increased GABA signaling due to receptor up-regulation has been shown to enhance motor performance^82^. It is important, however, to note that CB_1_ receptor levels rapidly increase during this exposure period in other brain areas related to motor function, including the basal ganglia, and cortex^53^.

However, it is also critical to remember that development requires a careful balance and timing of events, so changes in the rate of maturation of some developmental processes could have detrimental effects on behavior later in life^83^. Notably, cannabinoid exposure did not induce any long-term effects in motor coordination when tested on the parallel bar motor coordination paradigm during adolescence. Moreover, it is also possible that performance on the motor development task could be influenced by other performance variables (i.e. motivation).

In the current study, the effects of combined CP and ethanol exposure during development depended on the task. On the motor development tasks, performance of subjects exposed to the combination of CP and ethanol either showed initial delay (similar to ethanol exposure alone) followed by faster catch-up to the level of subjects exposed to CP only in the task, or they did not differ significantly from that controls. In other words, the opposing actions of the CP and EtOH were reflected in performance of subjects exposed to the combination of both drugs. Past studies have shown that cannabinoids can modulate ethanol’s teratogenic effects by protecting against ethanol-induced neurodegeneration or blocking ethanol’s effects^48,49,84,85^.

In contrast, the combination of CP and ethanol produced more severe motor coordination deficits than ethanol alone, at least among female subjects. This sex-specificity may have been due to a floor effect observed in the male subjects, as ethanol-exposed males performed very poorly. Nevertheless, these findings suggest that there may be synergistic effects of combined exposure and that offspring exposed to both drugs may be more severely impaired than if exposed to either drug alone.

Importantly, CP exposure elevated BACs, particularly among female subjects. This finding is consistent with both clinical^86^ and preclinical^87^ research suggesting that combined exposure to THC and alcohol leads to higher BACs compared to alcohol only. Higher BACs during pregnancy are associated with the severity of alcohol-induced brain related injuries^88^. In fact, this rise in BAC levels might help explain the high mortality rates observed in subjects exposed to both ethanol and cannabinoids. Higher BACs are also associated with more severe, long-lasting motor impairments as well^89,90^. Unfortunately, subjects who did not survive exposure typically died early in treatment (PD 4 or 5), before BAC data were collected (blood was collected on PD 6), so we do not have BAC data for these lost subjects.

The higher mortality following combined CP and EtOH exposure is notable. Prenatal alcohol exposure has been shown to increase mortality rates^91^, and although prenatal cannabinoid exposure may not increase mortality on its own^92^, one study has reported that combined exposure to prenatal cannabis and alcohol led to 100% fetotoxicity in mice and 73% fetotoxicity in rats, which is consistent with our elevated rates of mortality among the combination groups^93^. Moreover, the combination of cannabinoid and ethanol exposure during early postnatal development has been shown to be synergistically neurotoxic^48^. Unfortunately, mortality rates are not often reported, making it difficult to compare the mortality rates in this study with other studies. Nevertheless, given the increase in peak BAC caused by CP exposure, it will be important to determine whether CP may lead BACs to reach lethal levels.

We do recognize that the mortality rate in this group does mean that only a sub-population of subjects who underwent the combined drug exposure was tested on behaviors later in life; this could certainly have an impact on our outcome measures. However, it is most likely that those who survived and were tested on behavioral outcomes were likely the most resilient subjects. Despite this, we still found that combined exposure affected behavioral development. Thus, our results may underestimate the adverse consequences of combined exposure on behavioral development.

It is also important to note the general lack of dose response effects on most outcome measures among CP-exposed subjects in Experiment 1. Only the highest dose (CP4) yielded significant advancements in motor development throughout testing. However, there were no significant dose dependent effects on body growth or overall success on the motor development task. There are several possible explanations. First, the doses chosen for this study may have exceeded the asymptote of the dose response curve. Secondly, it is possible that the range of doses was not broad enough and captured a more static are of a curvilinear dose-response curve. Finally, as mentioned previously, the chosen exposure period (postnatal days 4-9) is a period of rapid growth in the cerebellum^94–96^ and in the endogenous cannabinoid system^52–55,97^, so variability in endogenous cannabinoid activity may have created more individual variation in response than differences due to dose (although the latter explanation is unlikely, given that the SEM was similar among CP-exposed and control subjects).

Past literature on behavioral effects following developmental CP exposure is very limited, and many studies examine effects of only a single dose. One study did show that a single administration of prenatal CP exposure induced ocular dysmorphology in a dose-dependent manner, although no behaviors were examined in that study^98^. Importantly though, eye defects were seen with doses as low as 0.0625mg/kg (range: 0.0625 – 2.0 mg/kg)^98^. Our dose range was much more limited (range: 0.1 – 0.4 mg/kg) and when comparing to the data on eye dsymorphology, the effects of dose on physical development within that range did not vary greatly. We exposed subjects to CP on multiple days, rather than a single dose, so given the variation in developmental age, duration of drug treatment, and differences in outcome measures, it is challenging to make direct comparisons.

Most past behavioral studies have focused on acute effects of single CP administration in adult rodents. The results of studies that have examined multiple doses vary greatly, depending on dose levels, ranges, administration route, and genetic strains. For example, one previous study showed learning impairments, but found no dose response differences, when administering CP i.p. within a range of 0.15 – 0.25 mg/kg, similar to the range used in the present study^99^. Others have reported dose-dependent behavioral effects with the same dose range that we used, but examined only 2 doses^59,100,101^. Again, acute effects in adults are not really comparable with the consequences of repeated exposures in neonates; nevertheless, even in some of those studies, they fail to see dose-dependent effects.

There are some important limitations of the present study to note. This study used CP, as it mimics the effects of developmental exposure to the primary psychoactive constituent of cannabis, THC. However, it is likely that effects may differ with THC itself, or the many variations of cannabis preparations. Over 80 cannabinoids have been identified in cannabis^102^, and some of them could potentially exacerbate or counteract the effects observed in this study. For example, cannabidiol has neuroprotective properties via anti-oxidative effects^103^, and cannabivarin, another cannabinoid, may be a weak cannabinoid receptor antagonist^104^. Moreover, the ratio of cannabinoids in cannabis preparations has varied widely over the years and across locations of preparation^17,18,105^. Future studies will need to consider the risk of prenatal exposure to various combinations of cannabinoids.

Lastly, this study specifically targeted exposure to cannabinoids and/or alcohol during part of the brain growth spurt (PD 4-9), a critical period of neural development^50,51^ involving the endogenous cannabinoid system^52^ and a period known to be sensitive to the effects of ethanol exposure^56,57^. However, it will be important to also examine drug exposure earlier in gestation and investigate how offspring’s behavioral development may vary depending on exposure period, as well as other routes of administration.

## 5. Conclusions

Cannabis is the most commonly used illicit drug among pregnant women, and approximately half of all women who report consuming cannabis during their pregnancy also report consuming alcohol, and likely do so simultaneously. This study indicates that not only can alcohol and cannabis exposure during the brain growth spurt affect motor development by themselves, but that combined exposure to alcohol and cannabis has unique effects, depending on outcome measure. Although cannabinoid exposure advanced motor development, countering ethanol-related delays, the combination led to more severe growth deficits and long-term impairments in motor coordination. Research that parallels the ongoing changes in prenatal cannabis and alcohol consumption trends has important implications for human 3^rd^ trimester fetal development, as combined exposure may be more damaging to a developing fetus than exposure to either drug alone. Given the recent legalization of cannabis in several areas of the world, elucidating the possible consequences of combined alcohol and cannabis consumption on brain and behavioral development is critical to guide future medical and public policy regarding drug use during pregnancy.

## Acknowledgements

Supported by NIAAA grant AA025425 to Dr. Thomas and an NIH Loan Repayment Program award to Dr. Breit. Special thanks to Dr. Nirelia Idrus, Cristina Rodriguez, and the Center for Behavioral Teratology at San Diego State University for assisting in data collection and interpretation.

